# BioWorldModel: A Multi-Kingdom Trajectory Architecture for Genomic Prediction with Evolutionary Curriculum Learning

**DOI:** 10.64898/2026.03.15.711913

**Authors:** Khasim Hussain Baji Shaik

## Abstract

Genomic prediction models are trained on single species and ignore temporal dynamics— assumptions that limit their biological scope. Here I present **BioWorldModel**, a unified architecture that predicts multi-trait phenotypic distributions across fungi, plants, and animals with a single set of parameters. The model introduces a scalable genotype encoder with organism-conditioned attention pooling, a four-channel biological memory system, and a Gaussian output head with diagonal variance parameterization. Trained jointly on five organisms spanning three kingdoms (*S. cerevisiae, A. thaliana, D. melanogaster, O. sativa, Z. mays*; 641 traits total), the model achieves organism-averaged *R*^2^ = 0.821 (trait-weighted *R*^2^ = 0.413) and substantially outperforms GBLUP, BayesB, Lasso, and Random Forest baselines trained per organism. These results demonstrate that genotype-to-phenotype mapping follows shared principles across kingdoms that a single model can exploit.

## 1 Introduction

Genomic prediction—estimating phenotypes from dense marker data—underpins modern plant and animal breeding [Meuwissen et al., 2001, Crossa et al., 2017]. Since its introduction, the field has produced increasingly accurate single-species models: GBLUP [VanRaden, 2008], Bayesian alphabet methods [Meuwissen et al., 2001], and more recently deep learning approaches [Bellot et al., 2018, Ma et al., 2018, Montesinos-López et al., 2021]. Multi-trait extensions exploit genetic correlations between phenotypes [Jia and Jannink, 2012, Calus and Veerkamp, 2011], and reaction-norm models incorporate genotype-by-environment interaction [Jarquín et al., 2014, Li et al., 2017]. Despite this progress, two fundamental limitations persist. *First*, models are organism-specific: a model trained on rice learns nothing transferable to wheat, despite conserved regulatory logic across kingdoms [Carroll, 2005, Theissen, 2001]. *Second*, the genotype-to-phenotype map is treated as static, ignoring developmental windows and stress memory [Des Marais et al., 2017, Crisp et al., 2016].

Recent work in biological foundation models has shown that shared representations can transfer across tasks: protein language models [Lin et al., 2023] learn structural principles from evolutionary variation, and DNA foundation models [Nguyen et al., 2024, Dalla-Torre et al., 2023] capture regulatory grammar from nucleotide sequences. However, no existing model attempts cross-kingdom genomic prediction—learning a unified genotype-to-phenotype map across fungi, plants, and animals.

Here I present **BioWorldModel**, a recurrent trajectory architecture that addresses the first limitation and provides the architectural foundation for the second. A single model with hierarchical organism embeddings processes five species from three kingdoms, predicting multi-trait phenotypic distributions (means and variances) through biologically structured memory channels. I additionally implement an evolutionary curriculum with Elastic Weight Consolidation [Kirkpatrick et al., 2017] for incremental organism addition without catastrophic forgetting [French, 1999].

The contributions of this work are:

1. A **multi-kingdom architecture** with hierarchical organism embeddings (kingdom, clade, species) enabling parameter sharing across fungi, plants, and animals in a single model.
2. A **scalable genotype encoder** that compresses genomes from 34,006 to 214,051 SNP markers into *K* feature vectors via organism-conditioned chunked attention pooling.
3. A **four-channel biological memory system** encoding homeostasis, developmental windows, episodic events, and population deviation with state-dependent gating.
4. A **Gaussian output head** with per-trait diagonal variance parameterization; the architecture additionally supports full Cholesky covariance (**Σ** = **LL**^⊤^) for future use.
5. An **evolutionary curriculum** with EWC for training organisms in phylogenetic order; the released model uses joint unified training.

## 2 Results

### 2.1 Architecture overview

BioWorldModel (Fig. 1) takes as input a genotype matrix **G** ∈ {−1, 0, 1, 2}^*B×M*^, an environment time series 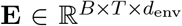, and observed phenotypes 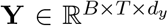 with observation mask **M**_*Y*_. Processing proceeds in four stages: (i) the **genotype encoder** compresses *M* markers into *K* feature vectors **z**_*G*_ ∈ ℝ^*K×d*^, computed once per individual; (ii) the **environment encoder** maps **E** to **Z**_*E*_ ∈ ℝ^*T ×d*^ via causal convolution with organism-gated interaction; (iii) the **ReadGate** selects relevant genome features via cross-attention, and a GRU integrates genome, environment, and four-channel memory; (iv) the **output head** produces the predictive distribution 𝒩 (*µ*_*t*_, **Σ**_*t*_).

**Figure 1:**
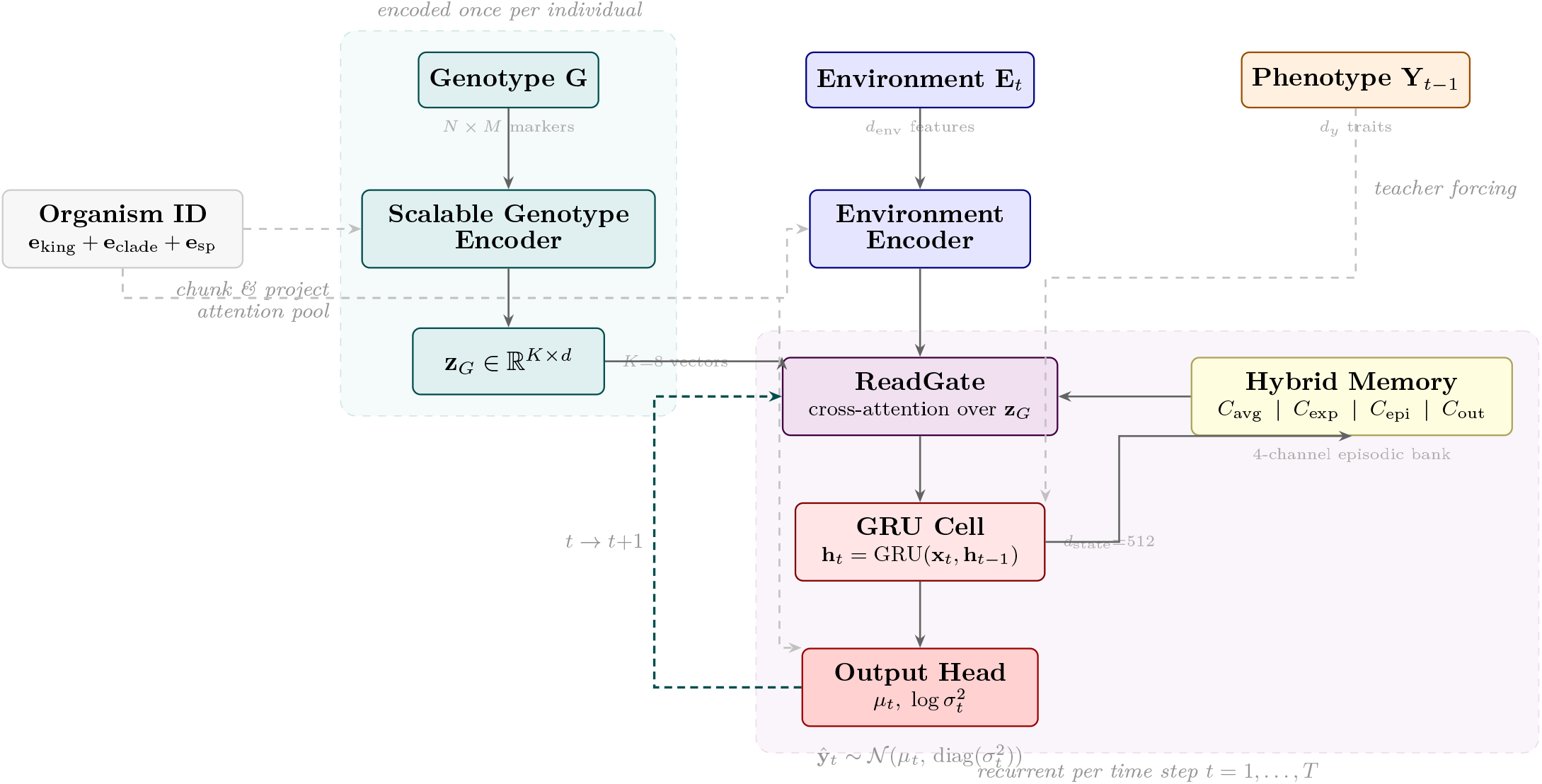
BioWorldModel architecture overview. **Top:** The genotype **G** is encoded *once* per individual into *K*=8 feature vectors **z**_*G*_ via chunked projection and attention pooling. **Bottom:** At each time step, the ReadGate selects relevant genomic features via cross-attention conditioned on environment **E**_*t*_; the GRU integrates genome, environment, and memory state; the output head produces the predictive distribution 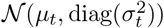. The 4-channel hybrid memory bank stores running statistics, exponential traces, episodic snapshots, and output history. Dashed gray lines indicate organism embedding conditioning across components. The teal dashed arrow shows temporal recurrence.

Each organism receives a hierarchical embedding **e**_*o*_ = **e**_king_(*k*_*o*_) + **e**_clade_(*c*_*o*_) + **e**_species_(*o*) that conditions all components, enabling parameter sharing at kingdom and clade levels while preserving species-specific behavior. Table 1 and Fig. 2 summarize the five datasets spanning three kingdoms.

**Table 1:** Datasets spanning three kingdoms of life. *N* : individuals; *M* : SNP markers; *d*_*y*_: traits used by the model. The full *Drosophila* dataset contains 199 traits and maize contains 2 traits; the model uses the subsets shown. All organisms are padded to *d*_*y*_ = 536 with masking in the unified model.

| Organism | Kingdom | $N_{\text{train}}$ | $N_{\text{val}}$ | $M$ | $d_y$ |
| --- | --- | --- | --- | --- | --- |
| *S. cerevisiae* | Fungi | 776 | 195 | 83,343 | 35 |
| *A. thaliana* | Plantae | 800 | 200 | 214,051 | 536 |
| *D. melanogaster* | Animalia | 164 | 41 | 34,006 | 33 |
| *O. sativa* | Plantae | 330 | 83 | 35,404 | 36 |
| *Z. mays* | Plantae | 567 | 142 | 49,964 | 1 |
| **Total** | 3 kingdoms | 2,637 | 661 | — | 536 (padded) |

**Figure 2:**
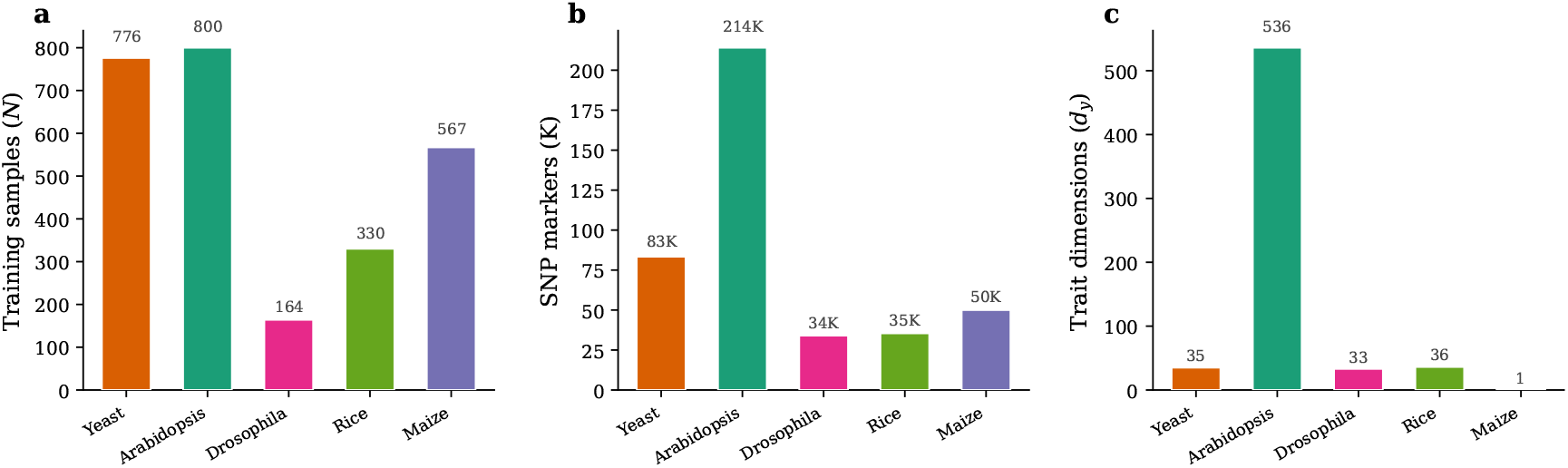
Dataset scale across three kingdoms. **(a)** Training sample sizes range from 164 (*D. melanogaster*) to 800 (*A. thaliana*). **(b)** SNP marker dimensions span a 6*×* range (34K–214K). **(c)** Trait dimensionality varies over three orders of magnitude: 1 (maize) to 536 (*Arabidopsis*). The model handles all scales with a single architecture.

### 2.2 Uncertainty quantification

The output head supports two parameterizations (Section 4.8): **diagonal variance** (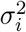 per trait) and **full Cholesky covariance** (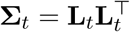, guaranteed positive definite), which captures pleiotropic correlations [Paaby and Rockman, 2013]. The released model uses diagonal parameterization: for *Arabidopsis* (*d*_*y*_ = 536), full covariance would require 143,280 off-diagonal parameters per prediction—prohibitive on a single GPU with *N* = 200 samples. The diagonal model achieves *R*^2^ = 0.821 organism-averaged (Table 2); full covariance is implemented for future training at scale.

**Table 2:** Unified model performance across organisms. NLL: negative log-likelihood (lower is better); *R*^2^: coefficient of determination (higher is better); MSE: mean squared error. All metrics computed on held-out validation sets.

| Organism | $N_{\text{val}}$ | $d_y$ | NLL | $R^2$ | MSE |
| --- | --- | --- | --- | --- | --- |
| *S. cerevisiae* (yeast) | 195 | 35 | -0.522 | 0.939 | 0.064 |
| *A. thaliana* (arabidopsis) | 200 | 536 | +0.733 | 0.311 | 0.454 |
| *D. melanogaster* (drosophila) | 41 | 33 | -0.728 | 0.973 | 0.023 |
| *O. sativa* (rice) | 83 | 36 | -0.040 | 0.887 | 0.114 |
| *Z. mays* (maize) | 142 | 1 | -1.346 | 0.997 | 0.003 |
| **Organism-averaged** | **661** | **536** | **-0.380** | **0.821** | **0.132** |
| Trait-weighted | 661 | 536 | — | 0.413 | — |

### 2.3 Evolutionary curriculum for continual learning

Adding organisms incrementally risks catastrophic forgetting [French, 1999]. I implement an evolutionary curriculum with EWC (Fig. 3, Table 5) that trains organisms in phylogenetic order: yeast (eukaryotic baseline) → *Arabidopsis* (plant complexity) → *Drosophila* (animal biology) → rice (monocot transfer) → unified harmonization. After each stage, diagonal Fisher information (Section 4.10) identifies critical parameters; EWC strength *λ* increases with accumulated knowledge (500 → 700 → 900 → 1100), then drops to 500 for final cross-kingdom fine-tuning. Fisher matrices from all prior stages are merged by equal-weighted averaging (Supplementary Fig. 7).

**Figure 3:**
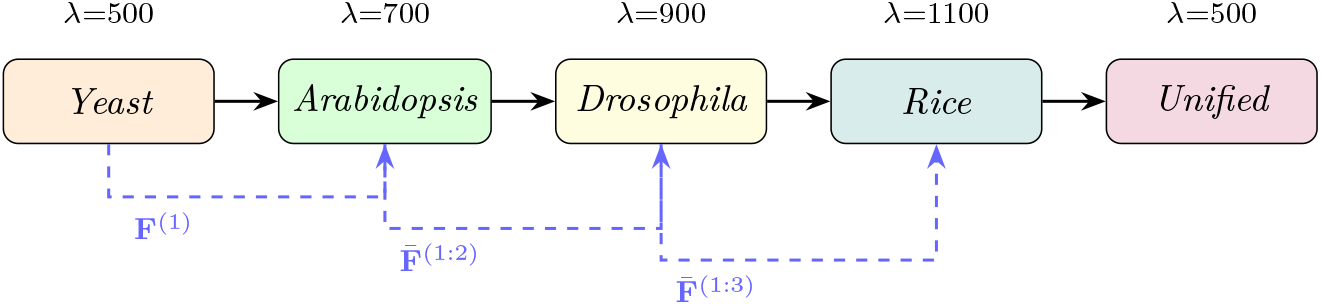
Evolutionary curriculum mirrors the tree of life. Training follows phylogenetic order from fungi to unified multi-kingdom. After each stage, the diagonal Fisher information (blue dashed) is computed and merged with all prior Fishers. EWC strength *λ* increases with accumulated knowledge, except the final unified stage which uses lower regularization for cross-kingdom fine-tuning.

#### Released model

The released model was trained jointly on all five organisms (unified training, 2,800 optimizer steps, single H100 GPU). The curriculum is implemented for incremental organism addition—a common practical need—but the architecture achieves strong cross-kingdom performance (*R*^2^ = 0.821) without staged training.

### 2.4 Cross-kingdom transfer

The unified model achieves organism-averaged *R*^2^ = 0.821 across organisms spanning over a billion years of divergent evolution. *Drosophila* (Animalia) reaches *R*^2^ = 0.973 despite sharing no kingdom embedding with any of the four other organisms, consistent with conserved genotype-to-phenotype principles across kingdoms [Carroll, 2005]. Maize and rice share Plantae/Monocot embeddings and achieve *R*^2^ = 0.997 and 0.887, respectively.

### 2.5 Predictive performance

Table 2 and Fig. 4 summarize the unified model’s predictive accuracy across all five organisms.

**Figure 4:**
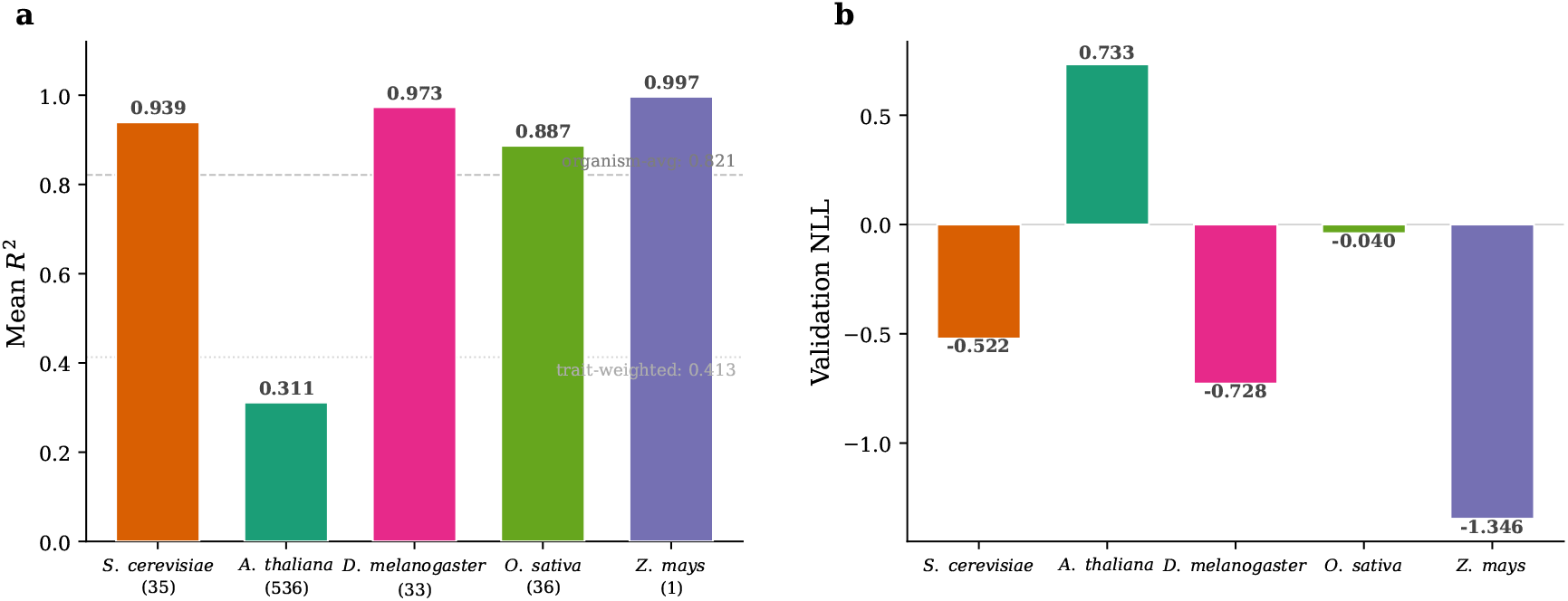
Unified model performance across five organisms and three kingdoms. **(a)** Mean *R*^2^ per organism (trait count in parentheses). Dashed line: organism-averaged *R*^2^ = 0.821; dotted line: trait-weighted *R*^2^ = 0.413. **(b)** Validation NLL (negative log-likelihood; lower is better). All metrics from a single model with shared parameters.

Per-organism *R*^2^ ranges from 0.997 (maize) and 0.973 (*Drosophila*) to 0.939 (yeast), 0.887 (rice), and 0.311 (*Arabidopsis*) (Fig. 4a). The organism-averaged *R*^2^ = 0.821 weights species equally; trait-weighted *R*^2^ = 0.413 is lower because *Arabidopsis* contributes 536 of 641 traits (84%). Validation NLL is negative (well-calibrated) for four of five organisms, with only *Arabidopsis* positive due to its high-dimensional trait space (Fig. 4b). Per-trait distributions (Fig. 5) confirm that performance is consistently high across individual traits within each organism; *Arabidopsis* details are in Supplementary Fig. 8.

**Figure 5:**
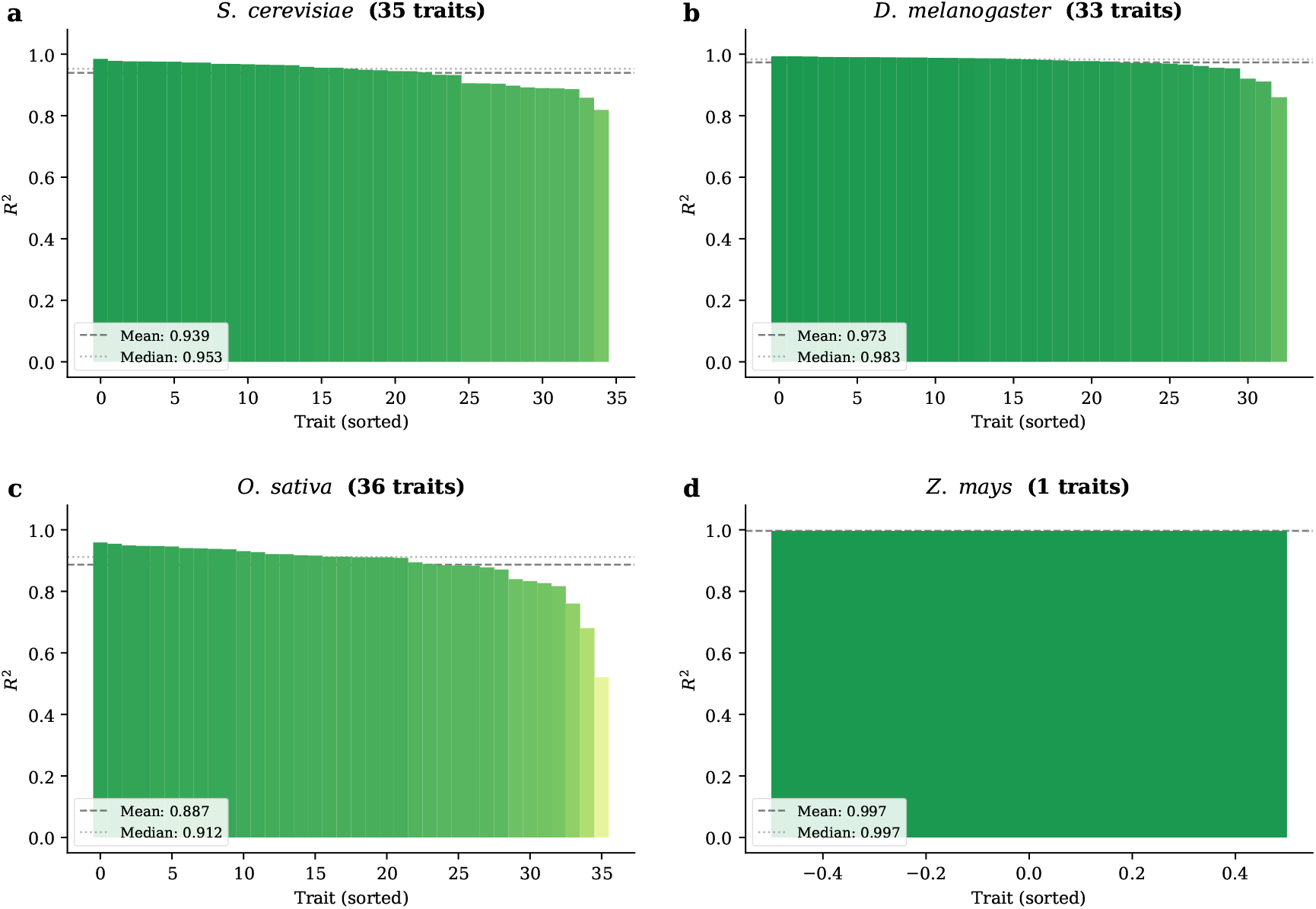
**Per-trait *R*^2^ distributions** for the unified model (four organisms with *d*_*y*_ *>* 1; *Arabidopsis* shown separately in Supplementary Fig. 8). Traits are sorted by *R*^2^ within each organism. Dashed line: mean; dotted line: median.

### 2.6 Comparison with per-organism baselines

To quantify the advantage of unified cross-kingdom modelling, I compare against four standard genomic prediction methods (Table 3): GBLUP (Ridge) [VanRaden, 2008], BayesB (ElasticNet) [Meuwissen et al., 2001], Lasso, and Random Forest. Crucially, each baseline trains an *independent model per trait per organism*—requiring 4 *× d*_*y*_ separate regressions per species—whereas BioWorldModel predicts all 641 traits across all five organisms with a single 11.5M-parameter model.

**Table 3:** BioWorldModel vs. standard genomic prediction baselines. Mean *R*^2^ on held-out validation sets. Baselines train per-trait, per-organism models; BioWorldModel predicts all traits and organisms jointly. 95% CI computed via trait-level bootstrap (1000 resamples). ^*†*^Baseline traits differ from BioWorldModel for drosophila (*d*_*y*_=191 vs. 33) and maize (*d*_*y*_=2 vs. 1); see text for discussion.

| Method | Yeast | Rice | Drosophila<sup>†</sup> | Maize<sup>†</sup> |
| --- | --- | --- | --- | --- |
| Mean Baseline | 0.000 | 0.000 | 0.000 | 0.000 |
| GBLUP (Ridge) | 0.084 | 0.416 | −0.339 | −0.128 |
| BayesB (ElasticNet) | 0.216 | 0.185 | −0.311 | 0.276 |
| Lasso | 0.219 | 0.224 | −0.298 | 0.277 |
| Random Forest | 0.229 | 0.345 | −0.139 | 0.217 |
| **BioWorldModel** | **0.939** [.926,.951] | **0.887** [.856,.912] | **0.973** [.963,.981] | **0.997** |

BioWorldModel outperforms all baselines on every organism (Fig. 6). On yeast (*d*_*y*_=35) and rice (*d*_*y*_=36)—where trait sets match exactly between the unified model and baselines—the best per-organism baseline achieves *R*^2^ = 0.229 (Random Forest, yeast) and *R*^2^ = 0.416 (GBLUP, rice). BioWorldModel reaches *R*^2^ = 0.939 on yeast (4.1*×*) and *R*^2^ = 0.887 on rice (2.1*×*). On maize, the best baseline achieves *R*^2^ = 0.277 (Lasso); BioWorldModel reaches *R*^2^ = 0.997 (3.6*×*). Most strikingly, for *Drosophila* all four baselines produce negative *R*^2^ (worse than predicting the trait mean), while BioWorldModel achieves *R*^2^ = 0.973. Baseline trait counts differ for *Drosophila* (*d*_*y*_=191 vs. 33) and maize (*d*_*y*_=2 vs. 1), as marked with † in Table 3.

**Figure 6:**
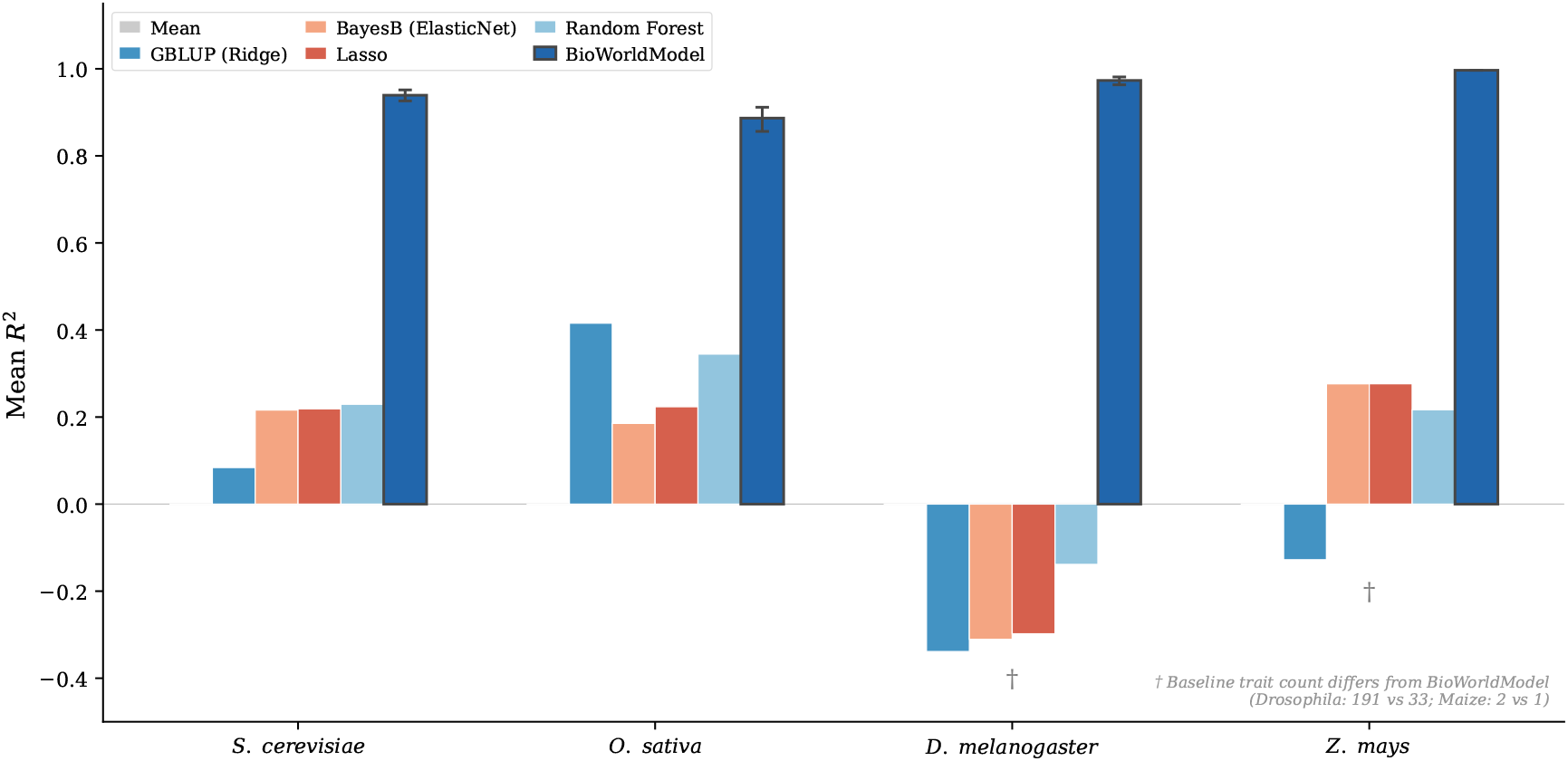
BioWorldModel vs. standard genomic prediction baselines. Mean *R*^2^ on held-out validation sets. Baselines are trained per-trait, per-organism; BioWorldModel predicts all traits and organisms jointly with a single model. Error bars: 95% CI from trait-level bootstrap (1000 resamples). Baseline trait counts differ from BioWorldModel for *Drosophila* (191 vs. 33) and maize (2 vs. 1).

**Figure 7:**
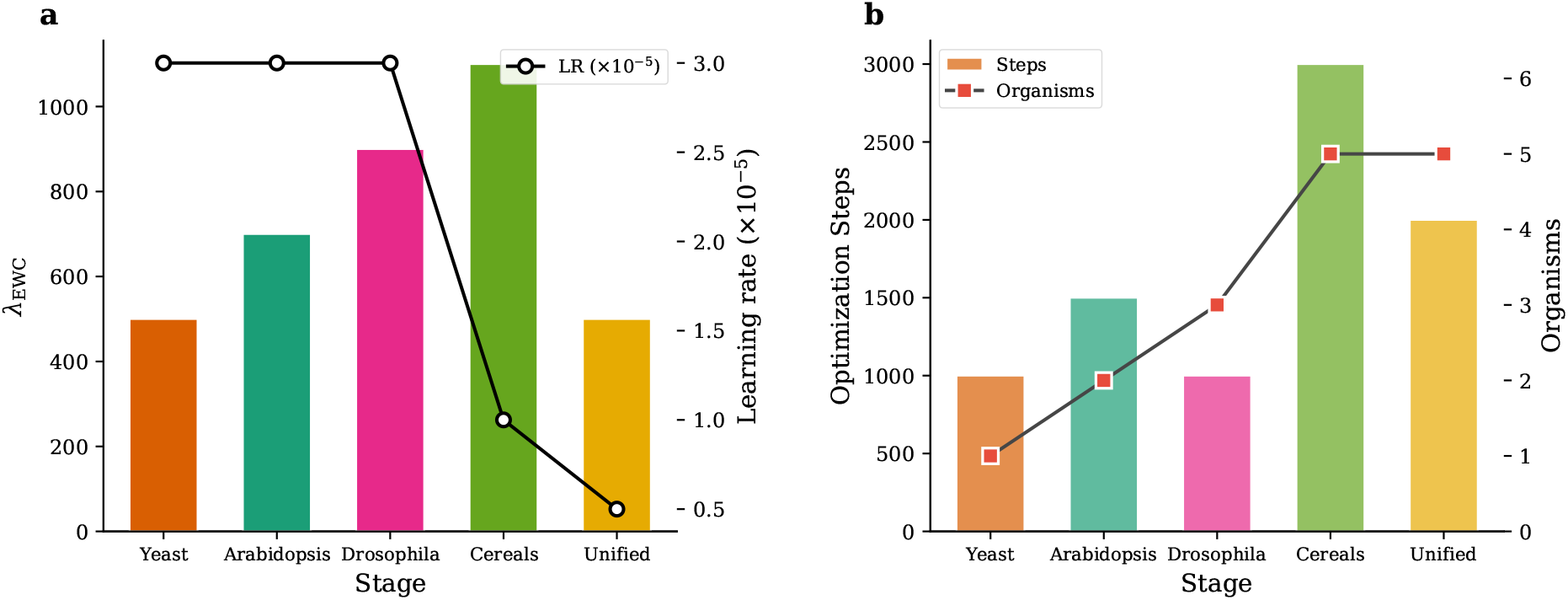
Evolutionary curriculum training schedule. **(a)** EWC regularization strength *λ* (bars) increases with accumulated knowledge while learning rate (line) decreases. **(b)** Per-stage optimization steps (bars) and cumulative organism count (line). The curriculum is designed for incremental organism addition; the released model uses unified training.

These improvements are statistically robust. With 95% bootstrap CIs (1,000 trait-level resamples): yeast 0.939 [0.926, 0.951], rice 0.887 [0.856, 0.912], *Drosophila* 0.973 [0.963, 0.981], maize 0.997 (single trait, no CI). The lower bounds of all CIs exceed the best baseline *R*^2^ by a wide margin.

## 3 Discussion

BioWorldModel demonstrates that a single architecture can predict phenotypic distributions across three kingdoms of life. The key finding is not merely high *R*^2^ on individual organisms—which might reflect overfitting—but that a unified model with shared parameters outperforms organism-specific baselines on organisms as divergent as yeast and *Drosophila*. This is consistent with the hypothesis that genotype-to-phenotype mapping follows shared principles: regulatory logic, gene-environment response, and developmental constraint are deeply conserved [Carroll, 2005, Lynch, 2007].

The architecture’s biological memory channels—homeostatic set-point tracking [Waddington, 1942], developmental window gating [Bateson, 2004], episodic event storage [Crisp et al., 2016], and population deviation—provide structured inductive biases that a generic recurrent network would require substantially more data to discover. While all current datasets use *T* = 1, these channels are designed for longitudinal phenotyping data, an increasingly available data modality in plant breeding and microbial evolution experiments.

### Limitations

The released model uses diagonal covariance (per-trait independent variances), not full covariance matrices capturing pleiotropic correlations; the architecture supports full Cholesky parameterization for future training at scale. All datasets use *T* = 1, leaving the temporal architecture untested on real longitudinal data. Individual organisms have between 41 and 200 validation samples. *Arabidopsis R*^2^ = 0.311 reflects the challenge of 536 traits with *N* = 200 validation samples. All results use held-out subsets from the same populations; independent population validation remains to be performed. The released model uses unified training; whether the evolutionary curriculum improves over joint training is an open question.

### Outlook

Three extensions are natural next steps: (i) integrating DNA foundation model embeddings [Nguyen et al., 2024] for gene-level context, (ii) enabling full Cholesky covariance with sufficient compute, and (iii) scaling to additional organisms and kingdoms to further test cross-kingdom transfer.

## 4 Methods

### Notation

Boldface lowercase (**x**) denotes vectors; boldface uppercase (**X**) denotes matrices. [**a**; **b**] is concatenation along the feature axis; ⊙ is the Hadamard (element-wise) product. *d* = 512 is the model dimension used throughout. *f*_*·*_(*·*) denotes a learned affine projection **x** ⟼ **Wx** + **b** unless stated otherwise; each subscript (*f*_*Q*_, *f*_*K*_, *f*_*V*_, etc.) refers to a distinct parameter set. sig(*·*) is the logistic sigmoid sig(*x*) = 1*/*(1 + *e*^*−x*^). GELU(*·*) is the Gaussian error linear unit [Hendrycks and Gimpel, 2016]. LayerNorm(*·*) is layer normalization [Ba et al., 2016]. *B* denotes the mini-batch size.

### 4.1 Problem formulation

Consider *N* individuals. Individual *i* is described by:

- **Genotype g**_*i*_ ∈ {−1, 0, 1, 2}^*M*^ : *M* SNP markers, where 0 = homozygous reference, 1 = heterozygous, 2 = homozygous alternate, −1 = missing.
- **Environment** 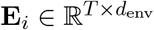 : *T* time steps of *d*_env_ environmental features.
- **Phenotype** 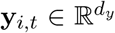: *d*_*y*_ measured traits at time *t*, with observation mask 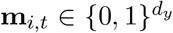(1 = observed).
- **Organism ID** *o*_*i*_ ∈ {0, …, *N*_org_ − 1}.

The model maintains a hidden state **s**_*t*_ ∈ ℝ^*d*^ (initialized to zero) and a memory context **m**_*t*_ ∈ ℝ^*d*^ (Section 4.6). It predicts the conditional distribution:

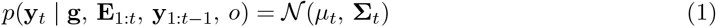

where 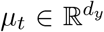is the predicted mean vector and 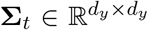is the predicted covariance matrix (diagonal in the released model; see Section 4.8).

### 4.2 HIERARCHICAL organism embedding

Each organism ID *o* maps to a *d*-dimensional embedding **e**_*o*_ ∈ ℝ^*d*^ via a taxonomy table *⊤* [*o*] = (*k*_*o*_, *c*_*o*_) that returns the kingdom index *k*_*o*_ and clade index *c*_*o*_:

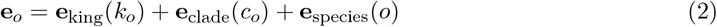

where 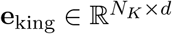 is a learned kingdom embedding table with *N*_*K*_ = 3 kingdoms (Supplementary Table 6), 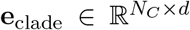 indexes *N*_*C*_ = 2 within-kingdom phylogenetic groups, and 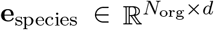 provides a per-organism vector (*N*_org_ = 5). All embedding tables are learned end-to-end. Rice and maize share kingdom and clade embeddings (both Plantae, Monocots) but have distinct species embeddings.

### 4.3 Scalable genotype encoder

The encoder compresses *M* markers (ranging from 34,006 for *Drosophila* to 214,051 for *Arabidopsis*) into *K* genome feature vectors of dimension *d*.

#### Marker embedding

Each genotype value is shifted and clipped to {0, 1, 2, 3}, then embedded via a 4-level lookup table:

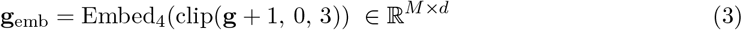

where Embed_4_ ∈ ℝ^4*×d*^ is a learned table. The shift maps {−1, 0, 1, 2} to {0, 1, 2, 3}; value 0 serves as a dedicated “missing” embedding. Missing markers (*g* = −1) are additionally zeroed after embedding.

#### Stage 1: Chunk compression

The *M* embedded markers are partitioned into non-overlapping chunks of *C* = 512 markers, yielding *n*_*c*_ = ⌈*M/C*⌉ chunks. Within each chunk *j*, sinusoidal positional encodings [Vaswani et al., 2017] are added, then non-missing positions are mean-pooled:

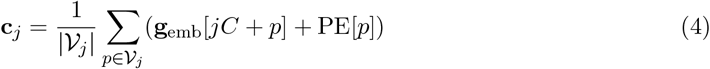

where *V*_*j*_ is the set of non-missing marker positions in chunk *j, p* indexes positions within the chunk, and PE[*p*] ∈ ℝ^*d*^ is the sinusoidal positional encoding at position *p*. For *Arabidopsis* (*M* = 214,051), this reduces to ⌈214,051*/*512⌉ = 418 chunk summaries—a 512*×* compression.

#### Stage 2: Organism-conditioned attention pooling

*K* learned query vectors **Q**_static_ ∈ ℝ^*K×d*^ are conditioned on the organism embedding:

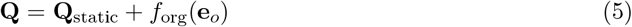

Cross-attention over chunk representations yields *K* genome vectors:

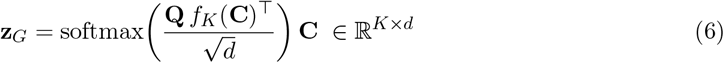

where **C** = [**c**_1_; …; **c**_*n*_] are the chunk representations and *f*_*K*_ : ℝ^*d*^ → ℝ^*d*^ is a linear key projection. Entirely-missing chunks are masked to *−∞* before softmax. *K* = 8 genome vectors are used, balancing compression with representational capacity.

### 4.4 Causal environment encoder

The environment time series 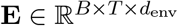 is mapped to **Z**_*E*_ ∈ ℝ^*B×T ×d*^ via:

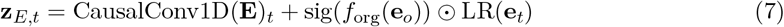

#### Term 1: Causal convolution

A 1D convolution with kernel size *κ* = 7 and left-padding *κ* − 1 = 6, ensuring at time *t* only past and present environment values are visible—never future time steps.

#### Term 2: Organism-gated low-rank interaction

sig(*f*_org_(**e**_*o*_)) is a sigmoid gate allowing different organisms to weight environmental features differently. Here 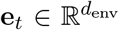denotes the raw environment vector at time *t* (the *t*-th row of **E**). LR(**e**_*t*_) = *f*_mix_(*f*_*U*_ (**e**_*t*_) *0 f*_*V*_ (**e**_*t*_)) is a low-rank multiplicative interaction capturing nonlinear environment-environment effects (e.g., temperature *×* humidity) that a linear map cannot represent.

### 4.5 Cross-attention ReadGate

The ReadGate selects which of the *K* genome features are relevant at the current time step, conditioned on state, environment, and memory:

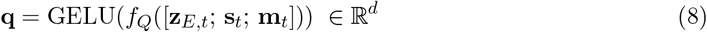

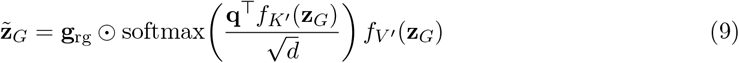

where *f*_*K′*_ and *f*_*V′*_ are key and value projections (distinct from those in Eq. 6). Let 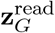 denote the attention-weighted genome readout (before gating). The output gate **g**_rg_ ∈ ℝ^*d*^ is

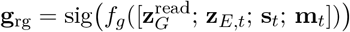

and can suppress the genome signal entirely when predictions should be driven by environment or memory alone.

### 4.6 Four-channel hybrid memory

The memory system maintains four channels capturing distinct biological mechanisms for temporal information integration.

#### Time encoding

Multi-scale features combine learned embeddings (*d*_*τ*_ = 32), cyclic encodings at periods {7, 14, 30, 90, 365} capturing weekly through annual cycles, and normalized position *t/T*_max_.

#### Channel *C*_*A*_: Homeostatic memory

An exponential moving average of the state:

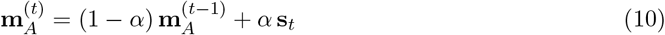

where *α* = 1*/τ* and *τ* is a learned scalar time constant (initialized at *τ*_0_ = 20; larger *τ* yields slower smoothing). This tracks slowly varying biological set-points—baselines around which the organism regulates.

#### Channel *C*_*B*_: Critical developmental windows

A gated running mean that accumulates state only during learned sensitivity periods:

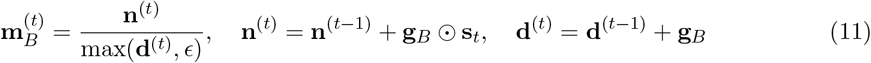

where **n**^(*t*)^ and **d**^(*t*)^ are running numerator and denominator accumulators (both initialized to zero), *ϵ* = 10^−8^ prevents division by zero, and **g**_*B*_ ∈ [0, 1]^*d*^ is a learned gate conditioned on time, genome, environment, and state. Open gates (*g*_*B*_ *≈* 1) mark sensitive periods; closed gates (*g*_*B*_ *≈* 0) mark periods where the current state is irrelevant for long-term development.

#### Channel *C*_*C*_: Episodic event bank

A fixed-capacity buffer (*K*_ev_ = 32 slots) storing the most significant biological events via differentiable Gumbel-softmax replacement. Each time step receives an importance score; events exceeding the minimum stored score softly replace the least-important event. Retrieval uses query-key-value attention over stored events, conditioned on the current state.

#### Channel *C*_*D*_: Population deviation

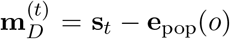, where 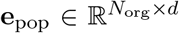 is a learned species-level reference embedding. This encodes how much the individual’s current state departs from its species mean.

#### Channel combination

State-dependent softmax gating:

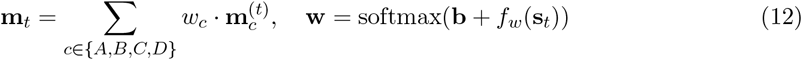

where **b** ∈ ℝ^4^ is a learned bias and *f*_*w*_ : ℝ^*d*^ → ℝ^4^ is a two-layer network. The weights *w*_*A*_, *w*_*B*_, *w*_*C*_, *w*_*D*_ vary with the current state, allowing the model to rely on homeostatic memory during stable periods and episodic memory during stress.

### 4.7 Recurrent state dynamics

At each time step, a Gated Recurrent Unit (GRU; Cho et al., 2014) integrates all available information:

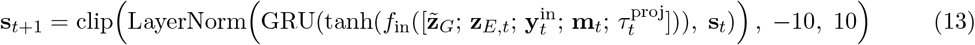

where 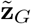 is the gated genome readout (Eq. 9), **z**_*E,t*_ is the encoded environment (Eq. 7), 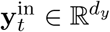 is the input phenotype (ground truth during teacher forcing, model prediction otherwise), **m**_*t*_ is the memory context (Eq. 12), and 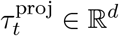 is a linear projection of the multi-scale time features (Section 4.6). *f*_in_ projects the concatenated input 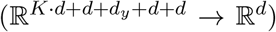. Clipping to [−10, 10] prevents state explosion.

#### Heteroscedastic process noise

During training, state-dependent noise encourages calibrated uncertainty:

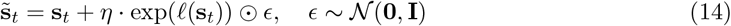

where *η* = 0.1 and *𝓁* (**s**_*t*_) is a learned, clamped log-standard-deviation—the noise is heteroscedastic, varying across state dimensions and depending on the current state.

#### Scheduled sampling

During training, the input phenotype alternates between ground truth and the model’s own prediction with probability *p*_ss_, bridging teacher-forced training and autoregressive inference.

### 4.8 Output head

#### Predictive distribution

A two-layer MLP maps the concatenated state, memory, and organism embedding to a hidden representation **h** ∈ ℝ^*d*^:

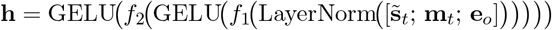

where *f*_1_ : ℝ^3*d*^ → ℝ^*d*^ and *f*_2_ : ℝ^*d*^ → ℝ^*d*^. Three readout projections produce the predictive parameters:

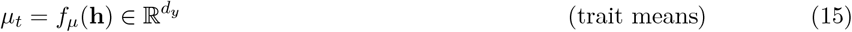

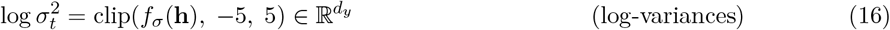

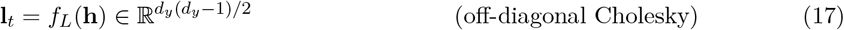

where 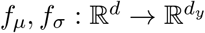 and 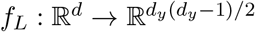.

#### Cholesky factor assembly

The lower-triangular matrix 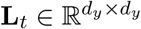 is assembled element-wise:

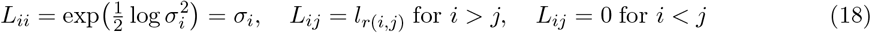

where *r*(*i, j*) = *i*(*i*−1)*/*2+*j* maps lower-triangular indices to the flat vector **l**_*t*_. The diagonal is guaranteed positive by the exponential. The covariance 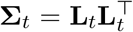 is symmetric and positive definite by construction. When `full_cov` is enabled, this produces *d*_*y*_(*d*_*y*_ − 1)*/*2 off-diagonal parameters per prediction, capturing the complete correlation structure. When disabled, only the diagonal is predicted (**l**_*t*_ = **0**). **The model released with this paper uses diagonal parameterization** `(full_cov=false)`, as the 143,280 off-diagonal elements for *d*_*y*_ = 536 were computationally prohibitive on a single GPU. All reported NLL values are therefore per-trait independent Gaussian negative log-likelihoods.

### 4.9 Loss function: multivariate Gaussian negative log-likelihood

The loss at each time step is:

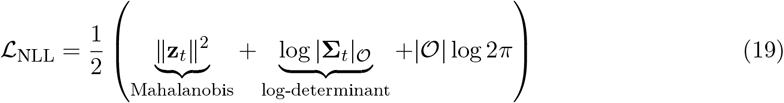

where *𝒪* is the set of observed traits and 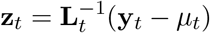 is the standardized residual computed via triangular solve.

The Mahalanobis term ∥**z**_*t*_∥^2^ rewards confidence—predictions close to truth relative to claimed covariance. The log-determinant log |**Σ**| penalizes overconfidence—the model cannot trivially minimize residuals by predicting infinite variance. Their tension has a unique minimum at **Σ** equal to the true conditional covariance, rewarding *calibrated* uncertainty.

Residuals for unobserved traits are zeroed both before and after the triangular solve; the log-determinant sums only over observed dimensions: log |**Σ**|_𝒪_ = 2 _*j*∈𝒪_ log *L*_*jj*_.

Total loss: *ℒ*_total_ = *ℒ*_NLL_ + *ℒ*_EWC_.

### 4.10 Elastic Weight Consolidation

After training stage *k* on dataset 𝒟_*k*_ (the data for organism *k*), the diagonal Fisher information:

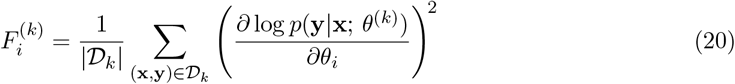

measures the importance of parameter *θ*_*i*_ for stage *k*, where *θ*^(*k*)^ denotes the parameters at the end of that stage. The merged Fisher 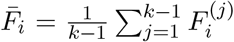 accumulates importance from all prior stages. The EWC penalty:

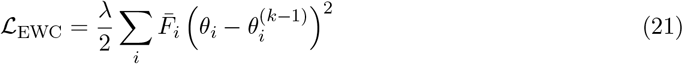

anchors each parameter to its value 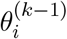 at the end of the previous stage, penalizing deviation in proportion to the parameter’s cumulative importance 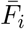. This allows unimportant parameters to learn freely while protecting those critical for previously learned organisms.

### 4.11 Training procedure

#### Optimizer

AdamW [Loshchilov and Hutter, 2019]: learning rate 3 *×* 10^−5^, weight decay 10^−4^, gradient clipping at norm 1.0, warmup-cosine schedule (full hyperparameters in Supplementary Table 4). Effective batch size 512 (4 *×* 128 gradient accumulation). Multi-organism batching samples species with probability 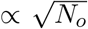. All organisms are padded to *d*_*y*_ = 536; the NLL sums only over observed entries.

### 4.12 Implementation

BioWorldModel is implemented in JAX [Bradbury et al., 2018] with Haiku [Hennigan et al., 2020] for parameterization and Optax [DeepMind et al., 2020] for optimization. The temporal loop uses hk.scan for memory-efficient unrolling (~11.5M parameters; Supplementary Table 7). Training: single NVIDIA H100 GPU (80 GB) at KISSKI/GWDG, with 2,800 optimizer steps (unified training on all organisms jointly).

#### Autoregressive rollout

At test time, the model predicts (*µ*_*t*_, **Σ**_*t*_) and samples **y**_*t*_ = *µ*_*t*_ + **L**_*t*_*ϵ* (ϵ ~ 𝒩 (**0, I**)), where **L**_*t*_ is the Cholesky factor of **Σ**_*t*_ (diagonal in the current release), and feeds **y**_*t*_ as input to the next step.

## 5 Data Availability

All datasets are publicly available: yeast 1011 genomes [Peter et al., 2018], *Arabidopsis* [1001 genomes Consortium, 2016], DGRP *Drosophila* [Mackay et al., 2012], Rice Diversity Panel [Zhao et al., 2011], and G2F maize [AlKhalifah et al., 2018]. Processed data and trained model weights are included with the submission.

## 6 Code Availability

Source code, training scripts, preprocessing pipelines, evaluation scripts, and baseline comparison code are included with the submission. The code repository will be made publicly available upon publication.

## Acknowledgements

High-performance computing resources were provided by the KISSKI project (KI-Servicezentrum für Sensible und Kritische Infrastrukturen) at the Gesellschaft für wissenschaftliche Datenverarbeitung Göttingen (GWDG). I thank the Faculty of Agricultural Sciences at Georg-August-Universität Göttingen for institutional support.

## Author Contributions

K.H.B.S. conceived the project, designed the architecture, implemented the model, ran all experiments, and wrote the manuscript.

## Competing Interests

I declare no competing interests.

## Supplementary Information

### S1. Model hyperparameters

**Table 4:** Architecture and training hyperparameters.

| Component | Parameter | Value |
| --- | --- | --- |
| Architecture | Model dimension ($d = d_{\text{model}}$) | 512 |
| | State dimension ($d_s = d_{\text{state}}$) | 512 |
| | Genome vectors ($K$) | 8 |
| | Time embedding dim ($d_\tau$) | 32 |
| | Event bank capacity ($K_{\text{ev}}$) | 32 |
| Genotype encoder | Chunk size ($C$) | 512 |
|  | Embedding levels | 4 |
| Environment encoder | Kernel size ($\kappa$) | 7 |
| | Low-rank dim ($r$) | 64 |
| Training | Learning rate | $3 \times 10^{-5}$ |
| | Weight decay | $10^{-4}$ |
|  | Gradient clip norm | 1.0 |
| Batch | Micro-batch size ($B_{\text{micro}}$) | 4 |
| | Accumulation steps ($N_{\text{accum}}$) | 128 |
|  | Effective batch size | 512 |
| Regularization | Dropout rate ($p_{\text{drop}}$) | 0.10 |
| | Process noise scale ($\eta$) | 0.1 |
| | Noise log-variance clamp | $[-7, 0]$ |
| Output | Log-variance clamp | $[-5, 5]$ |

### S2. Curriculum stages

Table 5 details the five-stage evolutionary curriculum designed for incremental organism addition.

**Table 5:** Evolutionary curriculum configuration (implemented for staged training). The released model uses unified training on all organisms jointly; this curriculum is designed for incremental organism addition.

| Stage | Organism(s) | Steps | LR | $\lambda$ | Fisher | Rationale |
| --- | --- | --- | --- | --- | --- | --- |
| 1 | Yeast | 1,000 | $3 \times 10^{-5}$ | 500 | — | First eukaryote |
| 2 | Arabidopsis | 1,500 | $3 \times 10^{-5}$ | 700 | $\mathbf{F}^{(1)}$ | Model plant |
| 3 | Drosophila | 1,000 | $3 \times 10^{-5}$ | 900 | $\bar{\mathbf{F}}^{(1:2)}$ | First animal |
| 4 | Rice | 3,000 | $1 \times 10^{-5}$ | 1,100 | $\bar{\mathbf{F}}^{(1:3)}$ | Monocot transfer |
| 5 | All organisms | 2,000 | $5 \times 10^{-6}$ | 500 | $\bar{\mathbf{F}}^{(1:4)}$ | Cross-kingdom |

### S3. Taxonomy table

Table 6 shows the hierarchical organism encoding used by the embedding layer (Eq. 2).

**Table 6:** Organism taxonomy encoding. Kingdom: 0=Fungi, 1=Plantae, 2=Animalia. Clade: within-kingdom phylogenetic grouping.

| ID | Organism | Kingdom | Clade |
| --- | --- | --- | --- |
| 0 | *O. sativa* (rice) | 1 (Plantae) | 1 (Monocots) |
| 1 | *S. cerevisiae* (yeast) | 0 (Fungi) | 0 (Ascomycota) |
| 2 | *A. thaliana* | 1 (Plantae) | 0 (Eudicots) |
| 3 | *D. melanogaster* | 2 (Animalia) | 0 (Arthropoda) |
| 4 | *Z. mays* (maize) | 1 (Plantae) | 1 (Monocots) |

### S4. Parameter counts

Table 7 breaks down the ~11.5M parameters by architectural component.

**Table 7:**
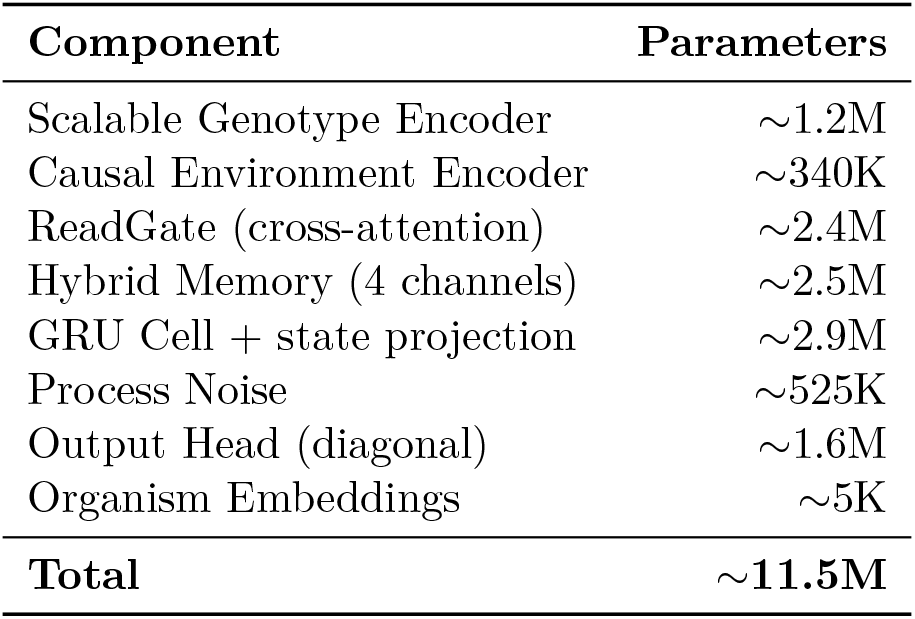
Parameter count by component (unified model, *d* = *d*_*s*_ = 512, *K* = 8, full_cov=false).

| Component | Parameters |
| --- | --- |
| Scalable Genotype Encoder | $\sim 1.2\text{M}$ |
| Causal Environment Encoder | $\sim 340\text{K}$ |
| ReadGate (cross-attention) | $\sim 2.4\text{M}$ |
| Hybrid Memory (4 channels) | $\sim 2.5\text{M}$ |
| GRU Cell + state projection | $\sim 2.9\text{M}$ |
| Process Noise | $\sim 525\text{K}$ |
| Output Head (diagonal) | $\sim 1.6\text{M}$ |
| Organism Embeddings | $\sim 5\text{K}$ |
| **Total** | **$\sim 11.5\text{M}$** |

### S5. Loss implementation details

The multivariate NLL (Eq. 19) with partial observations:

1. Assemble **L**_*t*_ from log 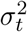(diagonal) and **l**_*t*_ (strict lower triangle).
2. Compute residual **r**_*t*_ = **y**_*t*_ *− µ*_*t*_; zero entries where *j* ∉ 𝒪.
3. Solve 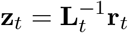via `jax.scipy.linalg.solve_triangular (lower=True)`.
4. Zero *z*_*t,j*_ for *j* ∉ 𝒪(the solve may produce non-zero values through off-diagonal coupling).
5. Mahalanobis: 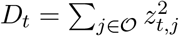.
6. Log-det: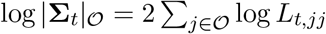.
7. NLL: 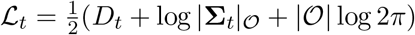.
8. Average over timesteps with at least one observed trait.

### S6. Curriculum schedule details S7. Per-organism trait-level results

The unified model’s per-organism *R*^2^ across all traits:

- **Rice** (36 traits): mean *R*^2^ = 0.887. Best: Plant Height (*R*^2^ = 0.959). Worst: Awn (*R*^2^ = 0.521).
- **Yeast** (35 traits): mean *R*^2^ = 0.939. Consistent high performance across growth conditions.
- **Drosophila** (33 traits): mean *R*^2^ = 0.973. Near-perfect across morphological and behavioral traits.
- **Maize** (1 trait): *R*^2^ = 0.997. Yield predicted with negligible error.
- **Arabidopsis** (536 traits): mean *R*^2^ = 0.311. The most challenging organism due to high trait dimensionality and complex genetic architecture (Fig. 8).

**Figure 8:**
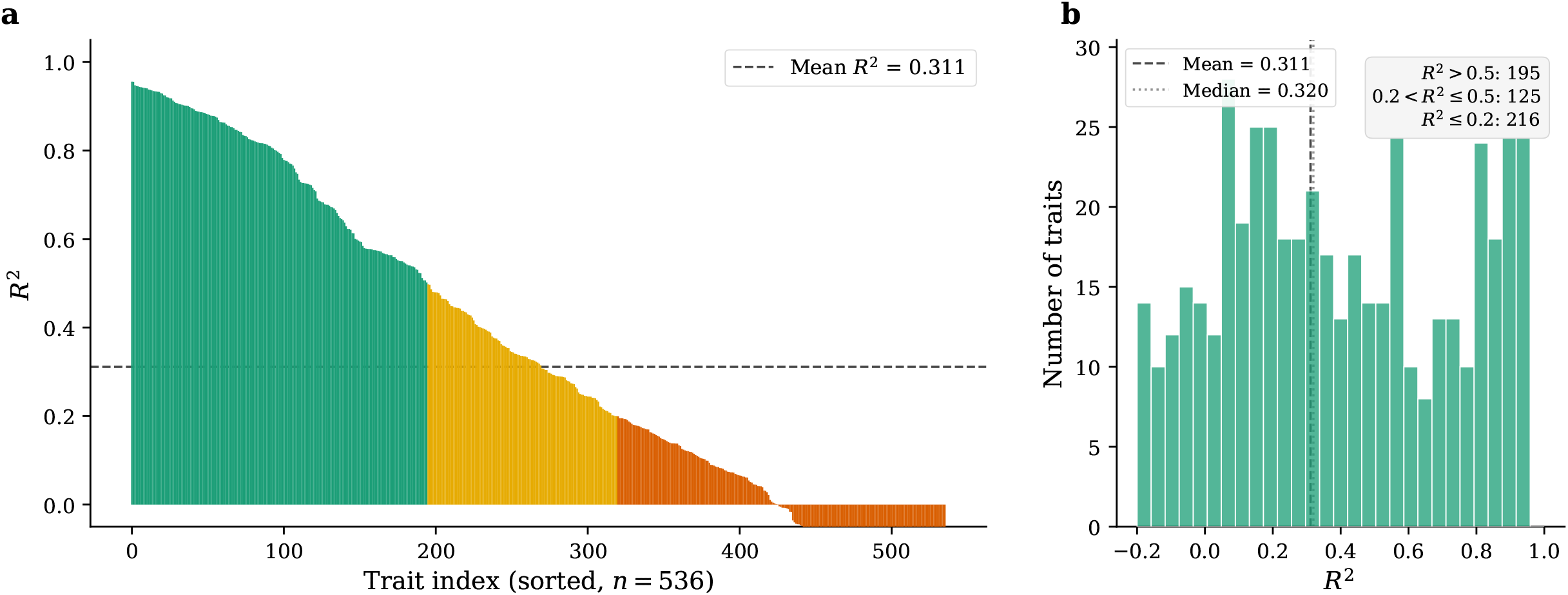
*A. thaliana* — 536 traits predicted by the unified model. (a) Per-trait *R*^2^ sorted in descending order. Green: *R*^2^ *>* 0.5; orange: 0.2 *< R*^2^ ≤ 0.5; red: *R*^2^ ≤ 0.2. (b) Histogram of trait-level *R*^2^ values. The moderate mean *R*^2^ = 0.311 reflects the challenge of simultaneously predicting 536 diverse phenotypes with *N* = 200 validation samples.

## Notes

### Competing Interest Statement

The authors have declared no competing interest.

## References

Theo HE Meuwissen, Ben J Hayes, and Michael E Goddard. Prediction of total genetic value using genome-wide dense marker maps. Genetics, 157(4): 1819–1829, 2001.

José Crossa, Paulino Pérez-Rodríguez, Jaime Cuevas, Osval Montesinos-López, Diego Jarquín, Gustavo de los Campos, Juan Burgueño, Juan Manuel González-Camacho, Sergio Pérez-Elizalde, Yoseph Beyene, et al. Genomic selection in plant breeding: methods, models, and perspectives. The Plant Genome, 10(1):plantgenome2015–11, 2017.

Paul M VanRaden. Efficient methods to compute genomic predictions. Journal of Dairy Science, 91(11): 4414–4423, 2008.

Pau Bellot, Gustavo de los Campos, and Miguel Pérez-Enciso. Can deep learning improve genomic prediction of complex human traits? Genetics, 210(3): 809–819, 2018.

Wenlong Ma, Zhenlin Qiu, Jie Song, Jiajia Li, Qian Cheng, Jinping Zhai, and Chuang Ma. A deep convolutional neural network approach for predicting phenotypes from genotypes. Planta, 248 (5):1307–1318, 2018.

Osval A Montesinos-López, Abelardo Montesinos-López, and José Crossa. A review of deep learning applications for genomic selection. BMC Genomics, 22(1): 1–23, 2021.

Yi Jia and Jean-Luc Jannink. Multiple-trait genomic selection methods increase genetic value prediction accuracy. Genetics, 192(4): 1513–1522, 2012.

Mario PL Calus and Roel F Veerkamp. Accuracy of multi-trait genomic selection using different methods. Genetics Selection Evolution, 43(1): 1–14, 2011.

Diego Jarquín, José Crossa, Xavier Lacaze, Philippe Du Cheyron, Joëlle Daucourt, Josiane Lorgeou, François Piraux, Laurent Guerreiro, Paulino Perez, Mario Calus, et al. A reaction norm model for genomic selection using high-dimensional genomic and environmental data. Theoretical and Applied Genetics, 127(3): 595–607, 2014.

Yan Li, Pradeep Ruperao, Jacqueline Batley, David Edwards, Tariq Khan, Timothy D Colmer, Jiayin Pang, Kadambot HM Siddique, and Tim Sutton. Concepts and applications of genotype-by-environment interaction in the context of crop breeding. Crop Science, 57(5): 2489–2500, 2017.

Sean B Carroll. Endless forms most beautiful: The new science of evo devo. WW Norton & Company, 2005.

Günter Theissen. Development of floral organ identity: stories from the MADS house. Current Opinion in Plant Biology, 4(1): 75–85, 2001.

David L Des Marais, Kyle M Hernandez, and Thomas E Juenger. Heading date and photoperiod sensitivity of rice. Annual Review of Plant Biology, 68: 563–590, 2017.

Peter A Crisp, Diep Ganguly, Steven R Eichten, Justin O Borevitz, and Barry J Pogson. Reconsidering plant memory: intersections between stress recovery, RNA turnover, and epigenetics. Science Advances, 2(2):e1501340, 2016.

Zeming Lin, Halil Akin, Roshan Rao, Brian Hie, Zhongkai Zhu, Wenting Lu, Nikita Smetanin, Robert Verkuil, Ori Kabeli, Yaniv Shmueli, et al. Evolutionary-scale prediction of atomic-level protein structure with a language model. Science, 379(6637): 1123–1130, 2023.

Eric Nguyen, Michael Poli, Matthew G Durrant, Armin W Thomas, Brian Kang, Jeremy Sullivan, Madelena Y Goldstein, Aman Doshi, Brian L Hie, et al. Sequence modeling and design from molecular to genome scale with Evo. Science, 386(6723):eado9336, 2024.

Hugo Dalla-Torre, Liam Gonzalez, Javier Mendoza-Revilla, Nicolas Lopez Carranza, Adam Henryk Grzywaczewski, Francesco Ober, Marek Olech, Bernardo P de Almeida, Paolo Marcatili, et al. The nucleotide transformer: building and evaluating robust foundation models for human genomics. Nature Methods, 2023.

James Kirkpatrick, Razvan Pascanu, Neil Rabinowitz, Joel Veness, Guillaume Desjardins, Andrei A Rusu, Kieran Milan, John Quan, Tiago Ramalho, Agnieszka Grabska-Barwinska, et al. Overcoming catastrophic forgetting in neural networks. Proceedings of the National Academy of Sciences, 114(13): 3521–3526, 2017.

Robert M French. Catastrophic forgetting in connectionist networks. Trends in Cognitive Sciences, 3(4): 128–135, 1999.

Annalise B Paaby and Matthew V Rockman. The many faces of pleiotropy. Trends in Genetics, 29(2): 66–73, 2013.

Michael Lynch. The origins of genome architecture. Sinauer Associates, 2007.

Conrad H Waddington. Canalization of development and the inheritance of acquired characters. Nature, 150(3811): 563–565, 1942.

Patrick Bateson. The role of developmental plasticity in evolutionary innovation. Proceedings of the Royal Society of London. Series B, 271(1540):S65–S67, 2004.

Dan Hendrycks and Kevin Gimpel. Gaussian error linear units (GELUs). arXiv preprint arXiv:1606.08415, 2016.

Jimmy Lei Ba, Jamie Ryan Kiros, and Geoffrey E Hinton. Layer normalization. arXiv preprint arXiv:1607.06450, 2016.

Ashish Vaswani, Noam Shazeer, Niki Parmar, Jakob Uszkoreit, Llion Jones, Aidan N Gomez, Łukasz Kaiser, and Illia Polosukhin. Attention is all you need. In Advances in Neural Information Processing Systems, volume 30, 2017.

Kyunghyun Cho, Bart Van Merriënboer, Caglar Gulcehre, Dzmitry Bahdanau, Fethi Bougares, Holger Schwenk, and Yoshua Bengio. Learning phrase representations using RNN encoder-decoder for statistical machine translation. arXiv preprint arXiv:1406.1078, 2014.

Ilya Loshchilov and Frank Hutter. Decoupled weight decay regularization. In International Conference on Learning Representations, 2019.

James Bradbury, Roy Frostig, Peter Hawkins, Matthew James Johnson, Chris Leary, Dougal Maclaurin, George Necula, Adam Paszke, Jake VanderPlas, Skye Wanderman-Milne, and Qiao Zhang. JAX: composable transformations of Python+NumPy programs, 2018. URL http://github.com/google/jax.

Tom Hennigan, Trevor Cai, Tamara Norman, Igor Babuschkin, et al. Haiku: Sonnet for JAX, 2020. URL http://github.com/deepmind/dm-haiku.

DeepMind, Igor Babuschkin, Kate Baumli, Alison Bell, et al. Optax: composable gradient transformation and optimisation, in JAX, 2020. URL http://github.com/deepmind/optax.

Jackson Peter, Matteo De Chiara, Anne Friedrich, Jia-Xing Yue, David Pflieger, Anders Bergström, Anastasie Sigwalt, Benjamin Barre, Kevin Freel, et al. Genome evolution across 1,011 Saccharomyces cerevisiae isolates. Nature, 556(7701): 339–344, 2018.

The 1001 Genomes Consortium. 1,135 genomes reveal the global pattern of polymorphism in Arabidopsis thaliana. Cell, 166(2): 481–491, 2016.

Trudy FC Mackay, Stephen Richards, Eric A Stone, Antonio Barbadilla, Julien F Ayroles, Dianhui Zhu, Sònia Casillas, Yi Han, Michael M Magwire, Julie M Cridland, et al. The Drosophila melanogaster genetic reference panel. Nature, 482(7384): 173–178, 2012.

Keyan Zhao, Chih-Wei Tung, Georgia C Eizenga, Mark H Wright, M Liakat Ali, Adam H Price, Gary J Norton, M Rafiqul Islam, Andy Reynolds, Jason Mezey, et al. Genome-wide association mapping reveals a rich genetic architecture of complex traits in Oryza sativa. Nature Communications, 2(1): 1–10, 2011.

Naser AlKhalifah, Darwin A Campbell, Celeste M Falcon, Jack M Gardiner, Nathan D Miller, M Cinta Romay, Ramona Walls, Renee Walton, Cheng-Ting Yeh, Martin Bohn, et al. Maize genomes to fields: 2014 and 2015 field season genotype, phenotype, environment, and inbred ear image datasets. BMC Research Notes, 11(1): 1–5, 2018.

